# Local and global crosstalk among heterochromatin marks drives epigenome patterning in Arabidopsis

**DOI:** 10.1101/2021.10.14.464341

**Authors:** Taiko Kim To, Chikae Yamasaki, Shoko Oda, Sayaka Tominaga, Akie Kobayashi, Yoshiaki Tarutani, Tetsuji Kakutani

## Abstract

Transposable elements (TEs) are robustly silenced by targeting of multiple epigenetic marks, but dynamics of crosstalk among these marks remains enigmatic. In Arabidopsis, TEs are silenced by cytosine methylation in both CpG and non-CpG contexts (mCG and mCH) and histone H3 lysine 9 methylation (H3K9me). While mCH and H3K9me are mutually dependent for their maintenance, mCG and mCH/H3K9me are independently maintained. Here we show that establishment, rather than maintenance, of mCH depends on mCG, accounting for the synergistic colocalization of these silent marks in TEs. When mCG is lost, establishment of mCH is abolished in TEs. mCG also guides mCH in active genes, although genic mCH/H3K9me is removed there. Unexpectedly, the targeting efficiency of mCH depends on relative, rather than absolute, levels of mCG, suggesting underlying global negative controls. We propose that the local positive feedback in heterochromatin dynamics, together with global negative feedback, drive robust and balanced epigenome patterning.

## Introduction

Large genomes of vertebrates and plants contain substantial amounts of transposable elements (TEs) and their derivatives. As TEs are a potential threat to genome stability and proper gene expression, they are silenced by epigenetic mechanisms such as cytosine methylation and histone H3 methylation at lysine 9 (H3K9me)^1–4^. In plant genomes, methylated cytosines are enriched in TEs for both CpG and non-CpG (or CpH, where H can be A, T, or C) contexts^5,6^. In Arabidopsis, methylation at CpG sites (mCG) is maintained by a DNA methyltransferase (MTase) called MET1 (METHYLTRANSFERASE 1)^7,8^. Methylation at CpH sites (mCH) is catalyzed by another class of DNA MTases, CHROMOMETHYLASE 2 and 3 (CMT2 and CMT3)^9–12^. These CMTs are recruited to regions with H3K9me. H3K9me is catalyzed by SUVH4, SUVH5 and SUVH6, and these H3K9 MTases, in turn, are recruited to regions with mCH^10,13,14^, generating a self-reinforcing positive feedback loop^15^. By this positive feedback, mCH and H3K9me are maintained through cell divisions.

While H3K9me and mCH depend on each other, their relationship to mCG has received less attention. Mutations in the CpG MTase gene *MET1* have only minor effects on mCH and H3K9me; similarly, mutations in CpH MTases CMTs or H3K9 MTases SUVHs also have only minor effects on mCG^5,6,13,16–18^. Thus, these two layers of modifications are maintained largely independently.

Although the two layers of modifications, mCG and mCH/H3K9me, are maintained independently, they are associated with each other during *de novo* establishment. This association in the establishment can be seen in both RNAi-dependent and -independent pathways. Plants can methylate both CpG and CpH sites in an RNAi-based pathway, called RdDM (RNA-directed DNA methylation). RdDM is a mechanism to trigger mCG and mCH by *de novo* DNA MTase DRM2, and its targeting depends on siRNAs and siRNA-associating RNAi components^19,20^. In addition to this well-investigated RNAi-based pathway, we have recently identified a very robust and precise RNAi-independent pathway to establish mCH and H3K9me *de novo* in coding regions of TEs (TE genes); both H3K9me and mCH are lost simultaneously in mutants for SUVHs or CMTs, but these marks recover efficiently and precisely in coding regions of TEs (TE genes) after reintroduction of the wild-type alleles^18^. Unexpectedly, this recovery of mCH/H3K9me is independent of the RNAi-based *de novo* DNA methylation machinery, such as DRM2 or RNA-dependent RNA polymerases. However, the recovery is inefficient in TE genes that lack mCG. These results suggest that mCG might induce *de novo* establishment of mCH and H3K9me in the RNAi-independent pathway.

Although both mCG and mCH/H3K9me are enriched in TEs, their distribution patterns differ in active genes. In addition to TEs, about 20% of active genes have mCG in their internal regions (gene bodies)^21–24^. In contrast, mCH and H3K9me are found almost exclusively in TEs^5,6^. A factor contributing to the exclusion of mCH/H3K9me from active genes is a Jumonji domain-containing histone demethylase gene, *INCREASE IN BONSAI METHYLATION 1* (*IBM1*)^25–27^; in *ibm1* mutants, H3K9me and mCH accumulate in expressed genes. Interestingly, the genic H3K9me and mCH in the *ibm1* mutant background are found in genes with body mCG^27^. Thus, mCG might direct *de novo* mCH in genes, as well as in TEs. However, a causative link between mCG and mCH/H3K9me remains to be examined in both genes and TEs.

Here, we directly examined whether mCG is necessary for the establishment of mCH by using a mutation of CpG MTase gene *MET1*. The results revealed that TE genes that lose mCG by the *met1* mutation failed to establish mCH. In addition to the effect on TE genes, the *met1*-induced loss of mCG compromised genic mCH accumulating in the background of *ibm1* mutant. Unexpectedly, the targeting efficiency of genic mCH depends on relative, rather than absolute, level of mCG. Furthermore, genome-wide accumulation of genic mCH caused a global decrease of mCH in TE genes, suggesting global negative feedback in the heterochromatin dynamics. Based on these and previous results, we propose that global negative feedback and local positive feedback of heterochromatin marks, combined with H3K9 demethylation in active genes, results in robust and balanced differentiation of silent and active genomic regions.

## Results

### Loss of MET1 function abolishes establishment of mCH in TE genes

Because mCH and H3K9me depend on each other, both modifications are lost both in the *cmt2 cmt3* double mutant of the mCH MTase genes (hereafter referred to as *cc*) and in the *suvh4 suvh5 suvh6* triple mutant of the H3K9 MTase genes (hereafter referred to as *sss*). The F1 progeny between these mutants inherited genomes without H3K9me and mCH, but all these mutated genes (*CMT*s and *SUVH*s) are complemented by functional wild-type alleles in heterozygous states in the F1 (Fig. 1a) ^18^. Within mCH, symmetric mCHG and asymmetric mCHH are often analyzed separately, because their controls are different; mCHG is catalyzed by CMT3 and CMT2, while mCHH is catalyzed by CMT2^11,12^. In the F1 progeny, mCH and H3K9me recovers efficiently despite their absence in the parents (Fig. 1a,c,i) ^18^. This efficient recovery is independent of RNAi, but inefficient in TE genes that lack mCG at transcription start site (TSS) ^18^.

**Fig. 1.**
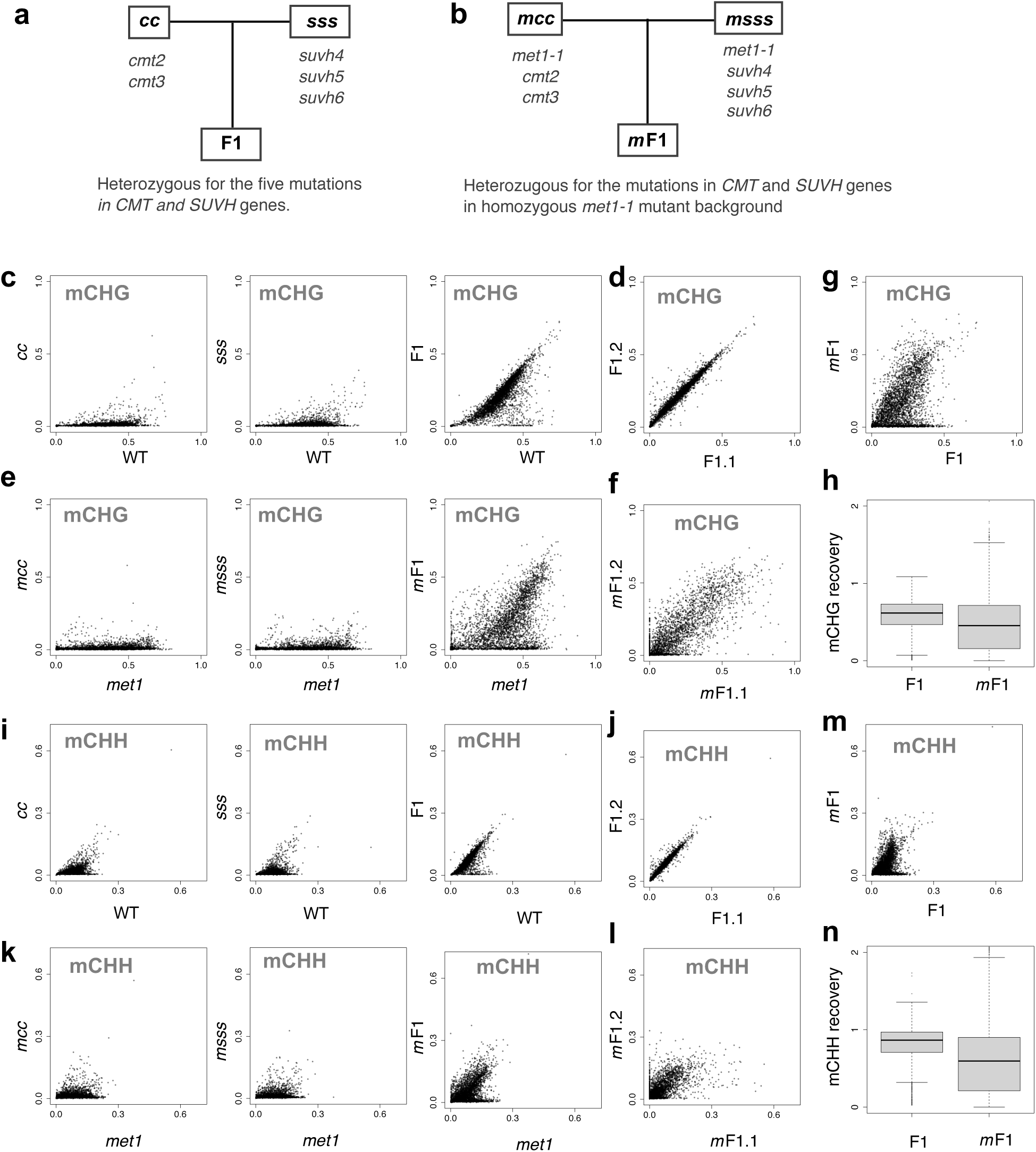
Loss of MET1 function abolishes establishment of mCH in TE genes. **a, b** Scheme of genetic crosses. Mutant of CpH MTases (*cmt2 cmt3* : *cc*) and mutant of H3K9 MTases (*suvh4 suvh5 suvh6* : *sss*) were crossed to generate F1, which are heterozygous for all the mutated genes (**a**). Analogous cross was done between F0 of *met1-1* mutant background (*mcc* and *msss*) to generate F1 with homozygous *met1-1* mutation (*m*F1) (**b**). **c** mCHG recovery in the F1. mCHG level of each TE gene in the mutants, *cc* (left), *sss* (middle), and the F1 (right), compared to a wild-type plant (WT). The results are reproduction of published results (GSE148753^18^). **d** A comparison of mCHG between the two individual F1 plants. **e** mCHG recovery in the *m*F1. mCHG level of each TE gene in the mutants *mcc* (left), *msss* (middle), and the *m*F1 (right), compared to a *met1-1* mutant. **f** A comparison of mCHG between the two individual *m*F1 plants. **g** A comparison of mCHG level between the F1 and *m*F1 plants. **h** The efficiency of mCHG recovery in F1 and *m*F1 plants. The efficiency of recovery was calculated as F1 / WT and *m*F1 / *met1*, respectively. To avoid division by values near zero, TE genes with mCHG (>0.1) in both WT and *met1* mutant were used (n=3212). Outliers deviated from the panel are not shown (n=11). **i**–**m** mCHH recovery of TE genes in the F1 and *m*F1 in the format of (**c**)–(**g**), respectively. **n** The efficiency of mCHH recovery in F1 and *m*F1 plants. The efficiency of recovery was calculated as F1 / WT and *m*F1 / *met1*, respectively. To avoid division by values near zero, TE genes with mCHH (>0.03) in both WT and *met1* mutant were used in (**n**) (n=2915). Outliers deviated from the panel are not shown (n=40). The original data for WT, *cc, sss* and F1 are from GSE148753^18^.

To directly investigate the possible contribution of mCG to the establishment of mCH, we conducted the same genetic experiments in the mutant background of CpG methyltransferase gene *MET1* (Fig. 1b). We used the incomplete loss-of-function allele, *met1-1*^28^, because the null allele of *met1* mutants in combination with mCH mutants result in severe developmental defects and infertility^29^. We generated the *met1-1 cmt2 cmt3* and the *met1-1 suvh4 suvh5 suvh6* mutants (hereafter referred to as *mcc* and *msss*, respectively) and performed a genetic cross between them to obtain the F1 plants (hereafter we call them as *m*F1; Fig. 1b), which can be compared to the original F1 plants between *cc* and *sss* (Fig. 1a). In contrast to the efficient recovery of mCH in F1 in the wild-type *MET1* background (Fig. 1c,d,i,j), the *m*F1 plants showed severe defects in the restoration of mCH in TE genes for both CHG (Fig. 1e–h) and CHH (Fig. 1k–n)contexts.

### Loss of mCG is associated with loss of mCH recovery

Next, we examined if the failure of mCH recovery in the *m*F1 was associated with the loss of mCG. Because *met1*-*1* is an incomplete loss-of-function allele, mCG remains in many, although not all, of TE genes^28,30^ (Supplementary Fig. 1a). Likewise, some TE genes kept mCG in the *mcc, msss* and the *m*F1 (Supplementary Fig. 1a). Thus, we compared the residual mCG and the efficiency of mCH recovery in the *m*F1. Indeed, mCH recovery was found where mCG remains (Fig. 2a,b). The residual mCG of the TE genes in the *m*F1 associated proportionally with their mCH recovery (Fig. 2c,d). TE genes with loss of mCG did not recover mCH (Fig. 2d), further demonstrating contribution of mCG to mCH establishment.

**Fig. 2.**
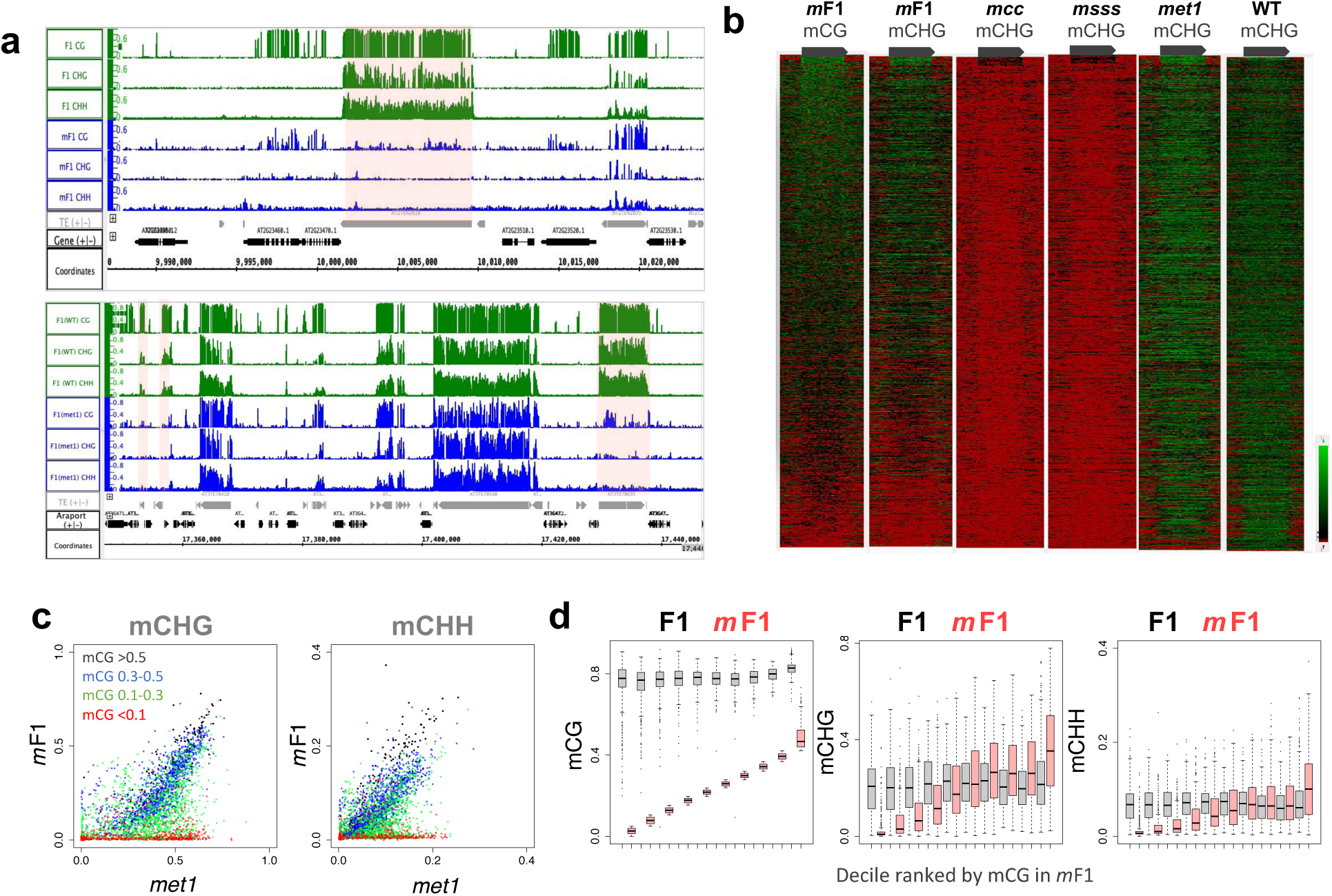
Loss of mCG is associated with loss of mCH recovery. **a** Genome Browser view of mCG, mCHG and mCHH in F1 and *m*F1 plants. TEs with loss of mCG fail to restore mCH in *m*F1 (highlighted in pink). The regions of Chr2/9,987,000-10,024,000 (upper) and Chr1/16,420,326-16,439,454 (lower) are shown. Grey and black arrows represent TEs and genes, respectively. **b** mCHG recovers in TE genes with remaining mCG. Each TE gene (length> 1000; n=2422) is aligned according to the mCG levels in *m*F1 and the methylation levels are shown in the form of heatmap for each genotype. Grey arrows represent TE genes. **c** Recovery of mCHG (left) and mCHH (right) is associated with the residual mCG in *m*F1. The scatter plots shown in the right panel of Fig. 1e and Fig. 1k were colored according to the residual mCG in *m*F1. Red : <0.1, green : 0.1–0.3, blue : 0.3–0.5, and black : >0.5 for mCG in *m*F1. **d** Boxplots showing the association between the residual mCG and mCH recovery in *m*F1. TE genes are divided into deciles according to the mCG levels in *m*F1, and their mCG (left), mCHG (middle) and mCHH (right) levels in F1 and *m*F1 plants are shown. TE genes with mCHG (>0.1) in both WT and *met1-1* were analyzed (n=3200). X axis represents mCG levels in *m*F1 divided into deciles and ordered from lower to higher. Each horizontal black line represents the median.

### Genic mCH is directed to regions with relatively high levels of mCG

Next, we asked if mCG in genes also affects their mCH. In wild-type plants, H3K9me2 and mCH are excluded from genes and that depends on the histone demethylase IBM1; in the *ibm1* mutant, H3K9me2 as well as mCH accumulate in genic regions^25–27^. Importantly, the genic mCH in the *ibm1* mutant correlates with the presence of mCG^26,27^. To see if mCG in gene bodies is responsible for the mCH in the *ibm1* background, we generated the double mutants of *met1-1* and *ibm1-4* (Fig. 3a, bottom left, *met1 ibm1*, hereafter referred to as *mi*) and examined the effect of the *met1* mutation on *ibm1*-induced genic mCH accumulation. In both *met1-1* and *mi* plants, genic mCG was drastically reduced, but they still had some residual mCG (Fig. 3b, top panel, Supplementary Fig. 1b and Supplementary Fig. 2). Unexpectedly, although the genic mCG is low in the *mi* mutant, a significant increase of genic mCH was detected (Fig. 3b, bottom panel and Supplementary Fig. 2). Nonetheless, the genic mCH in *mi* plants correlated with levels of their residual mCG (Fig. 3c–e), suggesting that the relative, rather than absolute, level of mCG is critical for the ectopic mCH in *mi* mutants. The spectrum of mCH and mCG in *mi* plants differed from that in *ibm1* plants and the spectrum differed even between different *mi* individuals (Supplementary Fig. 3a–c and Supplementary Fig. 4). Importantly, the genic mCH in each of *mi* plants correlated best to mCG level of the same plant but much less to those of the other plants (Fig. 3f and Supplementary Fig. 3d), suggesting that targeting of mCH is controlled by mCG, rather than other intrinsic properties of each gene.

**Fig. 3.**
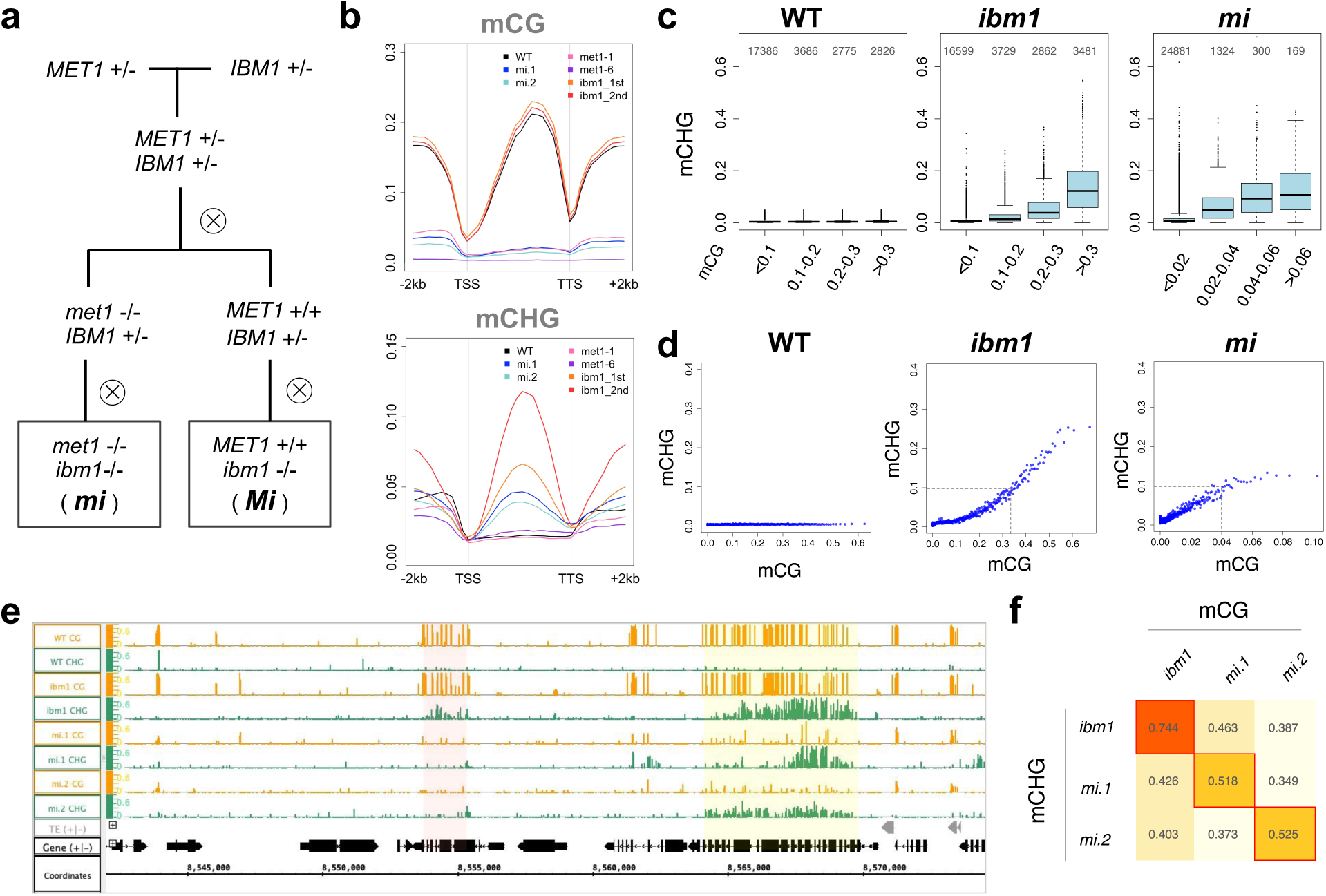
Genic mCH is directed to regions with relatively high levels of mCG. **a** Materials used. Starting from the cross between plants having *met1-1* (*met1*) or *ibm1-4* (*ibm1*) mutation in heterozygous state, the double heterozygote was self-pollinated to fix *met1* homozygote and *MET1* wild-type in the F2 generation. The *ibm1* mutation was fixed to be homozygous in the F3 generation. The F3 plants with homozygous *met1-1* mutation (*mi*) and homozygous wild-type *MET1* (*Mi*) were used in this study. **b** Averaged mCG and mCHG profiles over genes in a WT plant, *ibm1, met1 ibm1* (*mi*) and *met1* mutants. Values for *met1*-6, a null mutant of *MET1*, are shown for comparison (GSE148753). **c** mCHG levels are correlated with the mCG levels in both *ibm1* and *mi*. Genes are divided according to the mCG levels in the indicated genotypes and corresponding mCHG levels are shown by boxplots. The number of genes in each class is shown on the top. **d** The genes were sorted by the mCG levels of indicated genotypes, grouped in bins with 50 genes, and averaged mCHG levels in the bins are plotted against averaged mCG levels in the corresponding bins. Mean value is plotted for each bin. **e** Browser view of mCG (orange) and mCHG (green) in WT, *ibm1*, and two individual *mi* mutants. The region of Chr1/8,542,000-8,574,500 is shown. Grey and black arrows represent TEs and genes, respectively. Yellow shadow indicates an example of genic mCHG commonly seen in *ibm1* and *mi* mutants. It has residual mCG in the *mi* plants. Pink shadow indicates an example of genic mCHG seen in *ibm1* but not in *mi* mutants. **f** Correlation analysis of mCHG and mCG in the *ibm1* and two *mi* individual plants. The best correlated in the individuals are boxed in red. The number in the box represents the Pearson’s correlation coefficient. To exclude mis-annotated TEs, the genes with mCHG in the WT (>0.05) are excluded from the analysis in panels **b**–**d** and **f** (excluded n=1562; analyzed n=26723).

### Association of genic mCG and mCH in epigenetic variant individuals

mCH accumulated preferentially in genes with relatively high levels of mCG in *mi* as in *ibm1* single mutant plants. Unexpectedly, however, the level of mCG required for the accumulation of mCH seems much lower in *mi* than that in *ibm1*; 0.1 of mCHG was achieved when genes have about 0.04 of mCG level in *mi*, while more than 0.3 of mCG level in *ibm1* single mutant (Fig. 3d). One possible explanation for the difference is that it is attributable to the global reduction of mCG in *met1* background; in other words, the accumulation of genic mCH in *ibm1* mutant background may depend on the relative, rather than absolute, level of mCG. If the relative amount of mCG is really critical for the mCH, local loss of mCG is expected to show a more clear and strong effect. We next tested this hypothesis using materials with local and heritable loss of mCG, as stated below.

In plants, variation in mCG patterns tend to be transmitted stably over generations^31– 34^. That results in heritable loss of genic mCG detected among wild-type *MET1/MET1* siblings in the progeny of *MET1/met1* heterozygotes; and the spectrum of mCG loss is variable among individuals^35,36^. Consistent with these previous results, loss of genic mCG was detected in the *MET1/MET1 ibm1/ibm1* plants (hereafter referred to as *Mi*) originated from *MET1/met1-1 IBM1/ibm1* double-heterozygotes (Fig. 3a, bottom right and Supplementary Fig. 1c), and the spectrum of mCG differed among individual *Mi* plants (Fig. 4a). We used such heritable mCG variation among individuals to examine the association between local mCG loss and the ectopic mCH seen in the *ibm1* mutants.

**Fig. 4.**
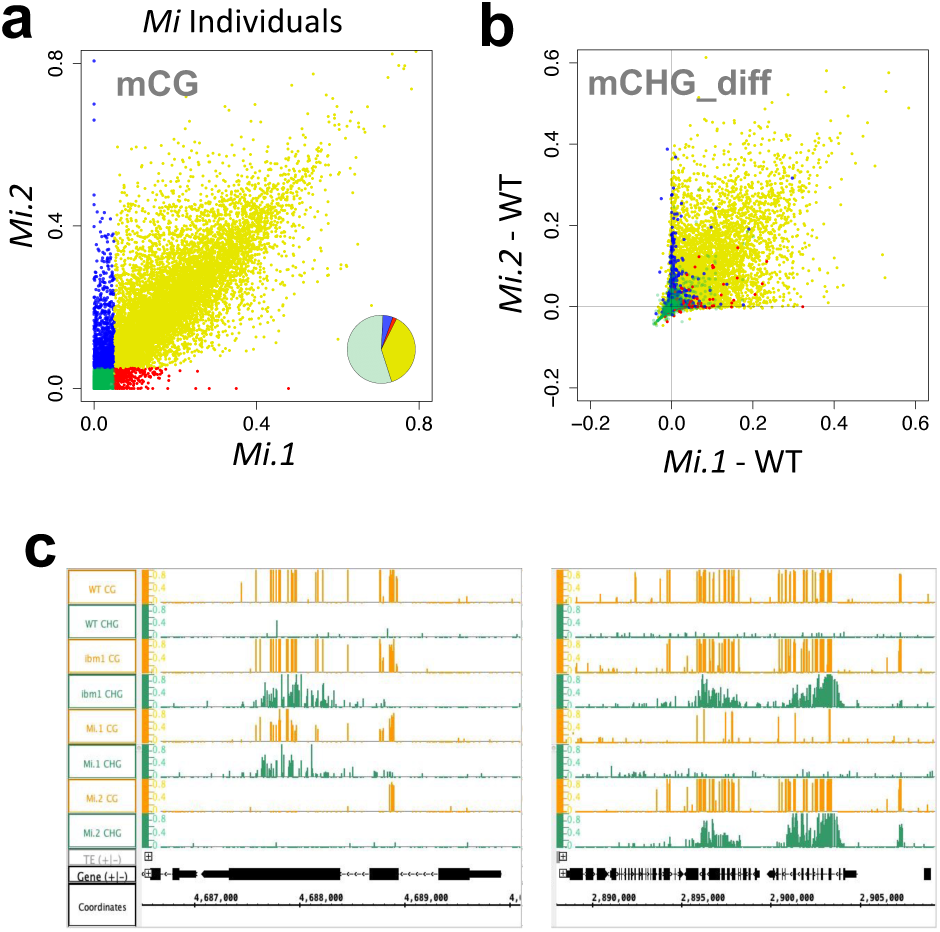
Association of genic mCG and mCH in epigenetic variant individuals. **a** Differential mCG levels of genes in the two *Mi* individuals. The genes with the mCG presence (>0.05) or absence (<0.05) in the individuals are colored in yellow (commonly CG methylated), green (commonly CG hypomethylated), red (CG methylated only in *Mi*.1 individual) and blue (CG methylated only in *Mi*.2 individual). The ratio of each group was shown as pie chart in the right bottom. **b** The increase in genic mCHG in the *Mi* individuals. The genes are colored according to the groups in (**a**). **c** Browser view of representative variable ectopic mCHG in *Mi* individuals. The gene in the left panel shows ectopic mCHG in *ibm1* and *Mi*.1 but not in *Mi*.2. The genes in the right panel show genic mCHG in *ibm1* and *Mi*.2 but not in *Mi*.1. The regions of Chr5/4,686,500-4,690,100 (left) and Chr1/2,888,000-2,906,200 (right) are shown. Grey and black arrows below represent TEs and genes, respectively. To exclude the mis-annotated TEs, the genes with mCHG in the WT (>0.05) are excluded from the analysis in (**a**,**b**) (excluded n=1562; analyzed n=26723).

In the two *Mi* siblings, genes with high mCG in only one of the two individuals (red and blue in Fig. 4a) show specific accumulation of mCH in that individual (Fig. 4b,c). In addition, genes with low mCG in both of the two individuals (green in Fig. 4a) show consistent lack of mCH accumulation (Fig. 4b). These results further demonstrate that genic mCH detected in the absence of *IBM1* function depends on mCG.

### mCH is controlled not only locally but also globally

The results above indicate that the relative, rather than the absolute, mCG level is critical for targeting of mCH seen in the *ibm1* mutant background. In other words, mCH/H3K9me is targeted to regions with mCG, but this targeting seems much enhanced when mCG levels are low in the other regions of the genome. The mCH levels may be controlled by a global negative feedback mechanism, which would explain the significant genic mCH in *mi* plants despite its low mCG.

In agreement with this interpretation, we have previously reported results suggesting global negative feedback to control heterochromatin marks, using another Arabidopsis mutant *ddm1* (*decrease in DNA methylation 1*)^11,37,38^. The *ddm1* mutation induces loss of mCG and mCH in heterochromatic TE genes^11^. The *ddm1*-induced loss of mC in TEs is associated with ectopic and stochastic gain of mC in other loci including genic regions^39,40^. Importantly, from a cross between *ddm1* and wild-type plants, the progenies in wild-type *DDM1* plants show the ectopic methylation even in regions originated from wild-type *DDM1* parent. This ectopic methylation correlates with amount of chromosome regions inherited from the *ddm1* parent, further demonstrating that loss of DNA methylation in TE genes induces genic mCH in *trans*^40^. The ectopic mCH is also induced in a *met1* mutant^32^, consistent with the idea that global loss of mCG accelerates targeting of mCH/H3K9me machinery via a global negative feedback mechanism.

The *ddm1* mutation abolishes both mCG and mCH. In addition to global mCG, global mCH levels may also affect mCH pattern formation by negative feedback. Gain of genic mCH in *ibm1* mutant induced reduction of mCH in TE genes (Fig. 5a). In the *ibm1* mutant, the genic mCH accumulate progressively over generations^40,41^, and the reduction of mCH in TE genes in *ibm1* was also progressive over generations (Fig. 5a), consistent with the idea that mCH levels are controlled globally by negative feedback. The genic mCH in *ibm1* mutation was much enhanced in the *ddm1 ibm1* double mutant (Fig. 5b,c), further suggesting that mCH levels are controlled by global negative feedback.

**Fig. 5.**
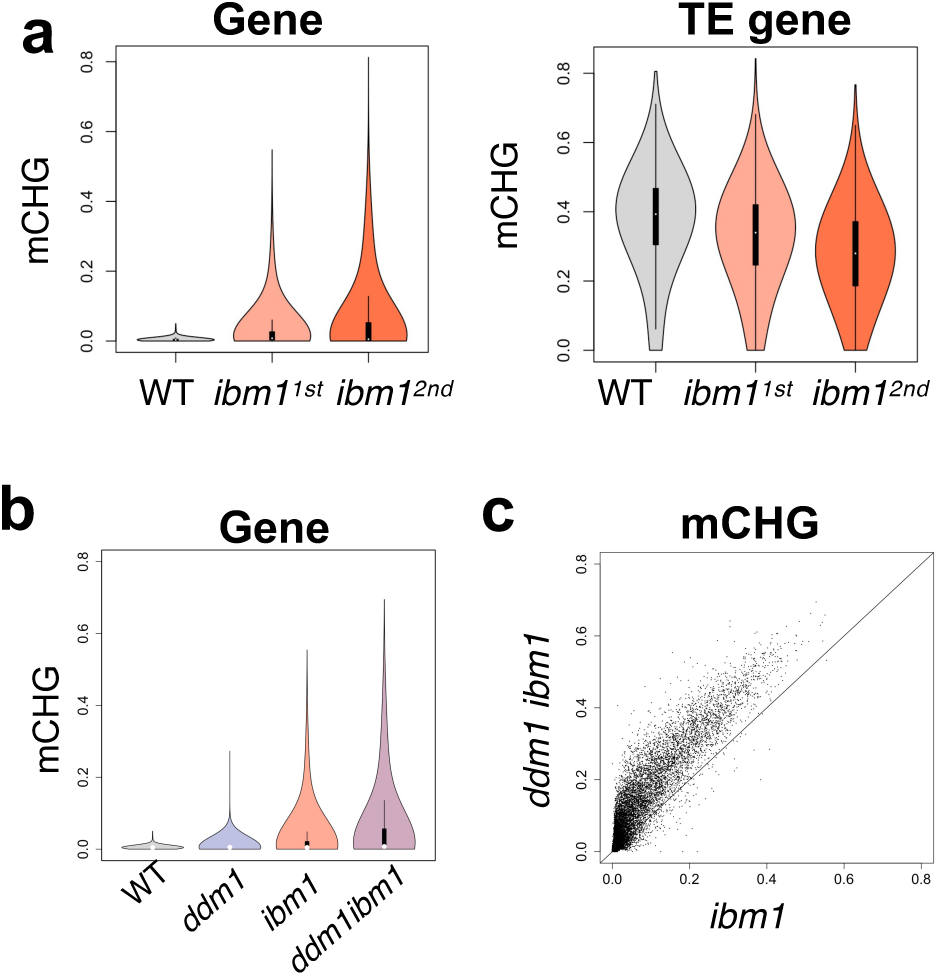
mCH is controlled not only locally but also globally. **a** Violin plots showing the progressive accumulation of mCHG in genes and decrease of mCHG in TE genes in the *ibm1* mutant. **b** Violin plot showing that *ddm1* mutation further enhanced the *ibm1*-induced accumulation of mCHG in genes. **c** The *ibm1*-induced genic mCHG was globally enhanced in *ddm1 ibm1* double mutant (line: y=x).

## Discussion

In plant genomes, both mCG and H3K9me2/mCH are important for silencing TEs, and both are enriched in TEs. Paradoxically, however, these two layers of modifications are maintained almost independently. Here we showed that mCG directs establishment of mCH, even though their maintenance is rather independent. Thus, we propose that colocalization of mCG and mCH in heterochromatin reflects this mechanistic link. Importantly, the very robust targeting of mCH we examined in this study is independent of RNAi for both TE genes^18^ and active genes^26^.

In contrast, RNAi is involved in targeting of mCG. The dynamics of mCG has been examined after its loss in *met1* mutation and reintroduction of a functional *MET1* gene^18,30,42^. In those systems, TE genes with efficient mCG recovery are associated not only with mCH but also with siRNAs, suggesting that RNAi and/or mCH also enhance establishment of mCG. When methylation is lost in both contexts in the *ddm1* mutants, the recovery is much slower than the cases in which methylation remains in one of the two contexts^18,31,43^. Thus, each of mCH and mCG facilitates establishment of the other.

RNAi-based mechanisms play especially significant part in non-coding regions of TEs^11,18^. siRNAs induce *de novo* DNA methylation by DRM2^19,20^. DRM2, in turn, also affects siRNA formation^44^. Furthermore, it was recently shown that siRNAs direct H3K9me in a DRM-independent pathway^45^. The emerging view would be that the three layers of marks (mCG, mCH/H3K9me, and siRNAs) interact locally to establish silent chromatin in both coding and non-coding regions (Fig. 6a). On the other hand, active chromatin states are stabilized by another positive feedback loop comprised of transcription and active H3K9 demethylation by IBM1 (Fig. 6a).

**Fig. 6.**
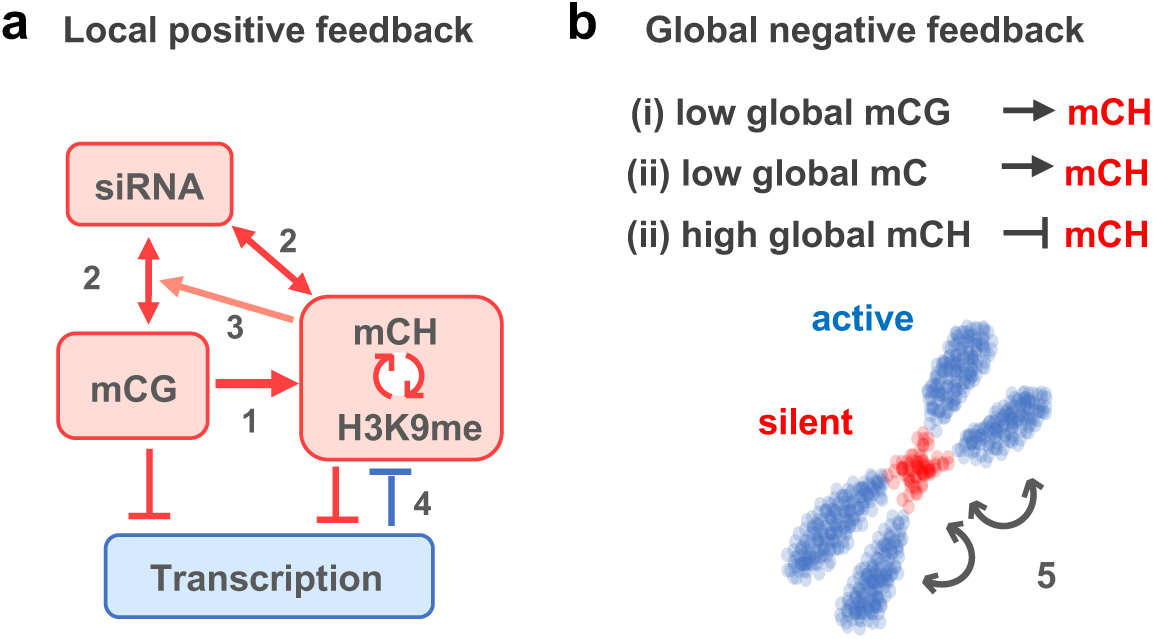
Local positive feedback and global negative feedback drive epigenome patterning. **a** Integrated model of local chromatin differentiation in Arabidopsis. (1) mCG directs precise and efficient *de novo* establishment of mCH/H3K9me in RNAi-independent pathway (This study and To et al. 2020^18^). (2) siRNA directs mCG and mCH by DRM2. In addition, DRM affects siRNA^44^. Furthermore, siRNA directs H3K9me in DRM2-independent pathway^45^. (3) mCH/H3K9me enhances siRNA-directed *de novo* mCG^18,30^. (4) Transcription induces demethylation of H3K9me by histone demethylase IBM1^27^. **b** A model for global negative feedback. In this model, we propose that the machinery to target mCH/H3K9me is negatively regulated by global level of mCG and mCH/H3K9me. The proposal is based on our three observations: (i) genic mCH is directed to regions with relatively high level of mCG, and this pathway is enhanced when global level of mCG is low (in *mi* in Fig. 3); (ii) when mCG and mCH of TEs decrease globally in the *ddm1* mutant, ectopic mCH is induced in genes^40^, and this increase is further enhanced in *ddm1 ibm1* mutant (Fig. 5 b,c); (iii) when genic mCH increases in the *ibm1* mutant, mCH in TE decreases (Fig. 5a). By this global negative feedback, amount of heterochromatin would not go to extreme, despite the strong local positive feedback mechanisms (**a**), resulting in balanced differentiation of active and silent chromosomal domains (5).

Although these multiple positive feedback mechanisms would stabilize and enhance silent and active states, positive feedback alone would have risk for the system to go out of control to excess. In addition to these local positive feedback mechanisms, our observation of *ibm1, met1*, and *ddm1* mutants revealed global negative feedback mechanisms to control genomic mCH level. Most significantly, global loss of mCG induces strong enhancement of machinery to direct mCH to regions with mCG (Fig. 3). In addition, mCH is also controlled negatively by global mCH level (Fig. 5a). Global negative feedback, combined with local positive feedback, would generate robust and balanced differentiation of active and silent genomic regions. The reaction-diffusion model is a powerful paradigm to understand pattern formation during development^46^, but that could also be applied to the pattern formation of active and inactive transcription units within the genome. The positive feedback mechanisms function locally to separate heterochromatic and euchromatic genomic domains, and the negative feedback by diffusible factor(s) controls the proportion of heterochromatic regions within the genome (Fig. 6b). Although the molecular basis for the global negative feedback remains unexplored, one possible mechanism could be a limitation of protein amounts involved in heterochromatin formation, such as CMTs and SUVHs.

Differentiation between genes and TE genes depends on histone demethylase IBM1 as well as on a chromatin remodeler DDM1; DDM1 is necessary for the TE-specific mCG, mCH and H3K9me^11,37,38,47^, but the underlying mechanisms have remained enigmatic. It has recently been shown that DDM1 binds to heterochromatin-specific H2A variant H2A.W and silence of TE genes in combination with H2A.W^48^. This pathway seems to be conserved in mammals; LSH1, the mammalian ortholog of DDM1, silences repetitive sequences by deposition of heterochromatin-specific H2A variant macro-H2A^49^. In Arabidopsis, another H2A variant, H2A.Z, negatively interacts with mCG^18,41,50,51^, and this pathway seems to be conserved to vertebrates^23^. Involvement of H2A variants in the local and global feedback mechanisms may also be an important target for future research.

## Methods

### Plant materials and growth conditions

The mutants *met1-1*^*28*^, *cc* (*cmt2 cmt3*) ^11^ and *sss* (*suvh4 suvh5 suvh6*)^52^ are kind gifts from Eric Richards, Daniel Zilberman and Judith Bender. The mutants *mcc* (*met1-1 cmt2 cmt3), msss* (*met1-1 suvh4 suvh5 suvh6), m*F1, *mi* (*met1-1 ibm1-4*), *Mi* (*MET1 ibm1-4*) and *ddm1 ibm1* were made in this study. The homozygous mutants of *met1-1 cmt2 cmt3* and *met1-1 suvh4 suvh5 suvh6* were used for genetic crosses to create the *m*F1 plants.

The plants were grown at 22 °C (16h light, 8h dark), firstly on MS agar media for 1– 2 weeks and then grown on soil. *ddm1 ibm1* plants were grown on MS agar media until harvesting.

### Whole Genome Bisulfite Sequencing and data processing

Genomic DNA was extracted from rosette leaves of one individual plant using the Nucleon Phytopure genomic DNA extraction kit (GE Healthcare), and whole genome bisulfite sequencing (WGBS) was performed as described previously^18^. Basically, two independent biological replicates were taken except for the wild-type Col-0, *met1-1* and the second generation of *ibm1-4* mutant. Genomic DNA was subjected to fragmentation using Focused Ultrasonicator (Covaris S220), and the size of 300–450bp were gel-extracted. The libraries were prepared using TruSeq DNA LT Sample Prep Kit (Illumina) and then subjected to bisulfite conversion using MethylCode Bisulfite Conversion Kit (Life Technologies). The resulting DNA were amplified using KAPA HiFi HotStart Uracil ReadyMix (Kapa Biosystems) and purified with Agencourt AMPure XP (Beckman Coulter). Raw sequence data and processed data were deposited in the GEO (GSE181896). The adaptor trimming and quality filtration were performed using Trimmomatic version 0.33^53^. The trimmed sequences were mapped to the Arabidopsis reference genome (TAIR10), deduplicated, and methylation data extracted using Bismark version 0.10.157^54^.

The annotations of genes and TEs are based on The Arabidopsis Information Resource^55^. The details of the annotation of TE genes is in TAIR website (https://www.arabidopsis.org). We used Perl scripts^18^ to count the numbers of methylated and total cytosines within a gene, and the methylation level for each context of each gene were calculated as the number of methylated cytosine within a gene divided by the number of total cytosine (weighted methylation level^56^). Rstudio (v1.1.463) was used to create scatter plots, box plots, and sliding window analysis. Genome Browser (Integrated Genome Browser^57^) was used to for browser views. To create the heatmaps, TE genes and their surrounding regions (2 kb) were divided into 20 and 10 segments respectively, then for each segment the value of methylated cytosines over total cytosines were calculated with Perl script. TE genes with the length (<1000, n=1038) were excluded. The processed data were visualized using TreeView 3^58^. The methylation recovery rates show in Fig. 1h and Fig. 1n, the methylation level of each gene in the F1 or *m*F1 were divided by that of corresponding background (WT or *met1*-1, respectively). To avoid division by values near zero, TE genes with low methylation level in WT and *met1-1* (CHG < 0.1 for Fig. 1h; n=318, or CHH < 0.03 for Fig. 1n; n=388) were excluded.

## Data availability

WGBS data in this study were deposited in the GEO with the accession number GSE181896. All other reasonable requests for data and research materials are available via contacting the corresponding author.

## Acknowledgements

We thank Eric Richards and Robert Schmitz for critical comments on the manuscript, and Judith Bender, Robert Fischer, Eric Richards and Daniel Zilberman for sharing mutant strains. Computations were partially performed on the NIG supercomputer at NIG, Japan. Supported by grants from, Japanese Ministry of Education, Culture, Sports, Science and Technology (26221105, 15H05963 19H00995 and 21H04977 to T.K., 19H05740 and 17K15059 to T.K.T.), CREST Grant, Japan (JPMJCR15O1 to T.K.), Systems Functional Genetics Project of the Transdisciplinary Research Integration Center, ROIS, Japan (to Y.T. and T.K.).

## Author Contributions

T.K.T., C.Y., S.O. and T.K. designed the study. T.K.T., C.Y., S.O., S.T., A.K., Y.T. and T.K. performed the experiments. T.K.T., C.Y. and S.O. analyzed the data. T.K.T. and T.K. wrote the paper with incorporating comments from the other authors.

