## Supplementary Figures for "Local and global crosstalk among heterochromatin marks drives epigenome patterning in Arabidopsis"

### Supplementary_Fig_1

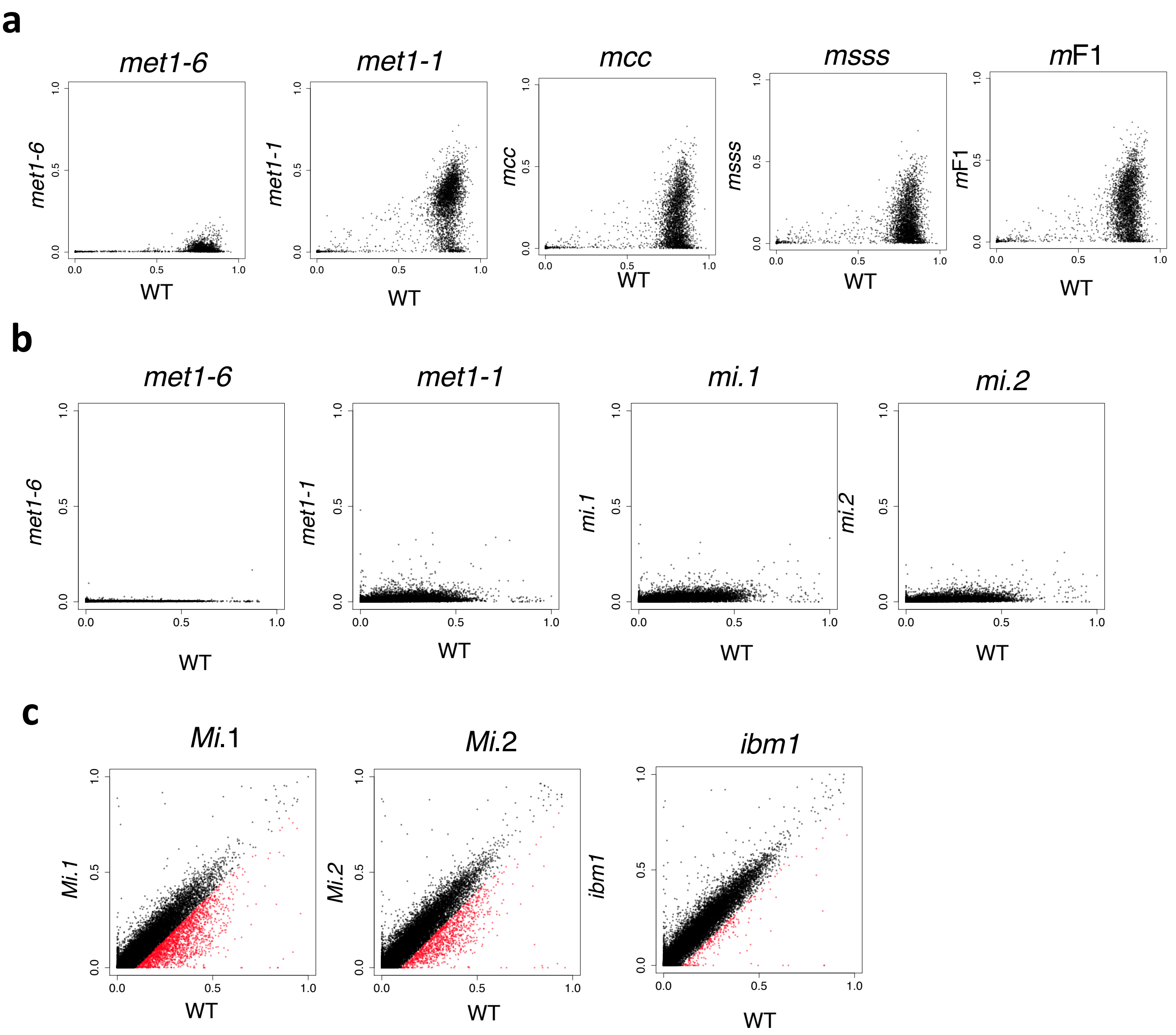


**Supplementary Fig. 1 Loss of mCG in the *met1* mutants.**

**a** The mCG levels for each TE genes in *met1-1* mutant compared to those of a wild-type (WT) plant. Results of *met1*-6, a null mutant of *MET1*, are shown in the left for comparison (GSE148753). **b** The mCG level compared between the *met1* mutants and WT for each of protein coding genes. **c** Loss of mCG detected in two individual *Mi* (*MET1/MET1 ibm1/ibm1* progeny originated from *MET1/met1-1 IBM1/ibm1* double heterozygote; as shown in Fig. 3a). mCG level of each protein coding genes in *Mi* plants compared to that in WT. Genes with mCG loss (WT - *Mi* >0.1) are shown in red. The *ibm1* mutant plant without experiencing *MET1* heterozygous state did not show such extensive loss of mCG (right panel).

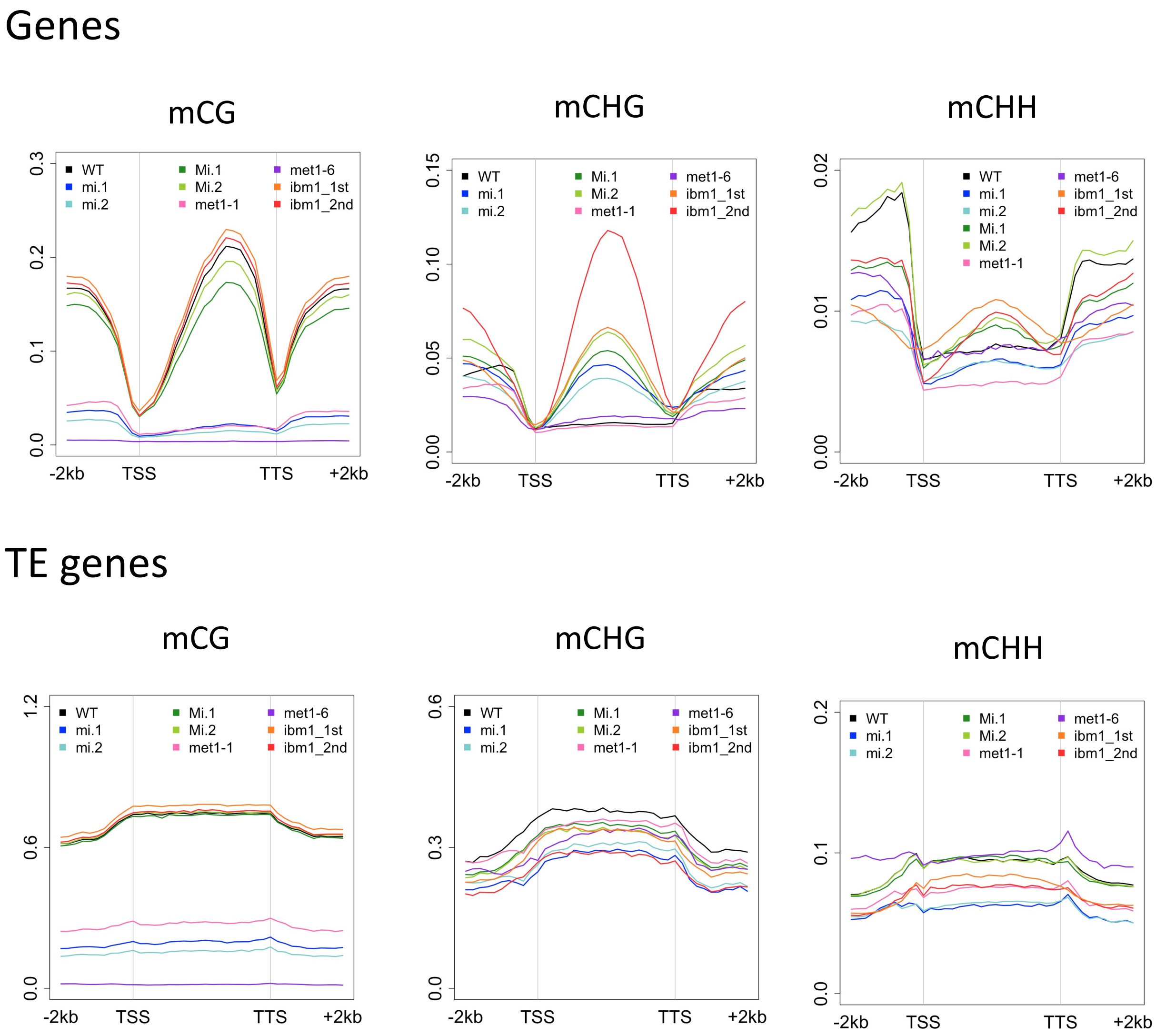


**Supplementary Fig. 2 mCG and mCH patterns over genes and TE genes.**

Averaged mCG (left), mCHG (middle) and mCHH (middle) over genes and TE genes. In each genotype, mean values are shown. To exclude the mis-annotated TEs, the genes with mCHG in the WT (>0.05) are excluded from the analysis for genes (excluded n=1562; analyzed n=26723).

### Supplementary_Fig_S3

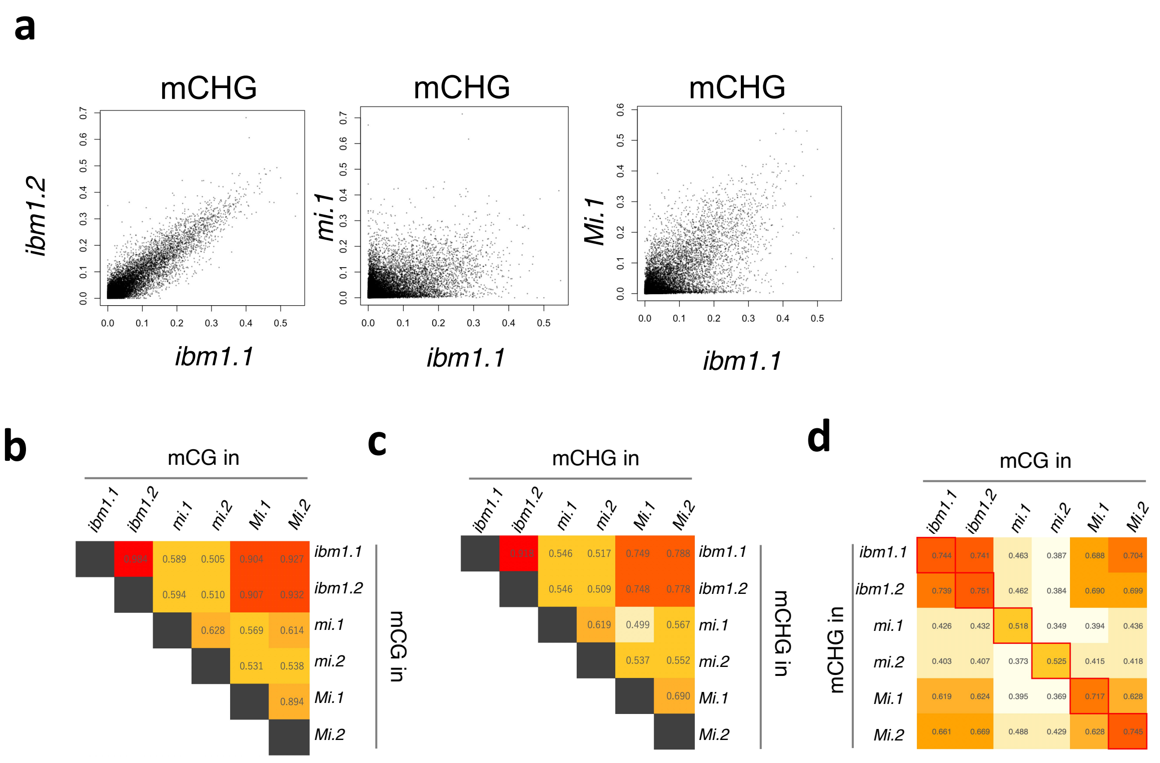


**Supplementary Fig. 3 The spectrum of genic mCH is different between *ibm1* and *mi*.** **a** Comparison of mCHG levels in the indicated genotypes. Spectrum of mCHG differs between *ibm1* and *mi*. **b** Correlation of mCG between the indicated individuals. The number in the box represents the pearson's correlation coefficient. **c** Correlation of mCHG between the indicated individuals. The number in the box represents the pearson's correlation coefficient. **d** Correlation between mCHG and mCG in the indicated individuals. The best correlated in the individuals are boxed in red. The number in the box represents the pearson's correlation coefficient. To exclude the mis-annotated TEs, the genes with mCHG in the WT (>0.05) are excluded from the analysis in panels (**a**)–(**d**) (excluded n=1562; analyzed n=26723).

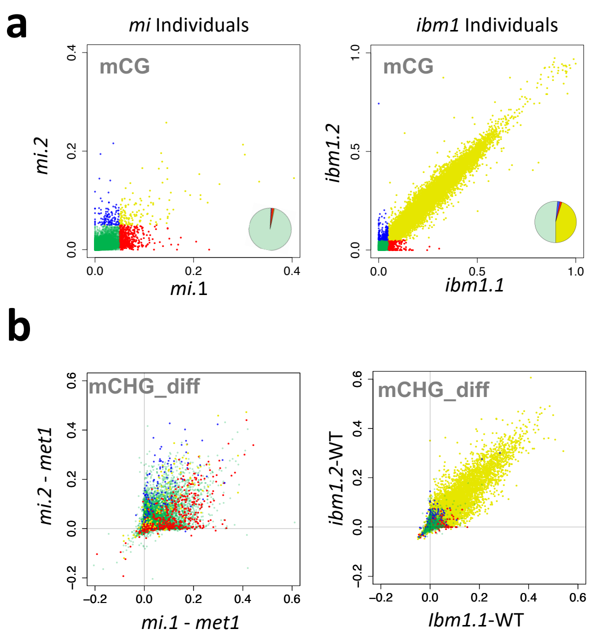


**Supplementary Fig. 4 Genic mCHG is correlated with the presence of mCG in the individual *mi* and *ibm1* plants.**

**a** Differential mCG levels of genes in the two *mi* and *ibm1* individuals. The genes with the mCG presence (>0.05) or absence (<0.05) in the individuals are colored yellow (commonly CG methylated), green (commonly CG hypomethylated), red (CG methylated only in *mi*.1 individual) and blue (CG methylated only in *mi*.2 individual). The ratio of each group was shown as pie chart. **b** The ectopic mCHG in genes in the *ibm1* individuals. The genes are colored according to the groups in (**a**). To exclude the mis-annotated TEs, the genes with mCHG in the WT (>0.05) are excluded from the analysis (excluded n=1562; analyzed n=26723).
